# HRWR: Predicting Potential Efficacious Drug Combination Based on Hypergraph Random Walk with Restart

**DOI:** 10.1101/2020.12.10.420760

**Authors:** Qi Wang, Guiying Yan

## Abstract

Some studies have shown that efficacious drug combination can increase the therapeutic effect, and decrease drug toxicity and side-effects. Thus, drug combinations have been widely used in the treatment of complex diseases, especially cancer. However, experiment-based methods are extremely costly in time and money. Computational models can greatly reduce the cost, but most of the models do not use the data of more than two drugs and lose a lot of useful information. Here, we used high-order drug combination information and developed a hypergraph random walk with restart model (HRWR) for efficacious drug combination prediction.

As a result, compared with the other methods by leave-one-out cross-validation (LOOCV), the Area Under Receiver Operating Characteristic Curve (AUROC) of the HRWR algorithm were higher than others. Moreover, the case studies of lung cancer, breast cancer, and colorectal cancer showed that HRWR had a powerful ability to predict potential efficacious combinations, which provides new prospects for cancer treatment. The code and dataset of HRWR are freely available at https://github.com/wangqi27/HRWR.

## Introduction

The drug combination is a fixed-dose combination, including two or more active drug components in a single dosage form, which has multiple advantages compared to monotherapy (Collier, 2012)(Liu et al., 2014): it offers higher efficacies or, through lower individual dosage, it can also decrease the drug side-effects and toxicity. Many studies have shown that synergistic drug combinations are widely used in the treatment of AIDs, cancer, and other complex diseases (Feliu et al., 2009). However, experimental screening efficacious drug combinations is time-consuming, costly, laborious, and inefficient. Thus, we are far from exploring a large number of possible combinations that could have potentially positive clinical effects. The mathematical model can produce a predictive result, we can regard the combination with a high prediction score as a potential efficacious drug combination. Small-scale drug trials can greatly reduce investment in human and material resources.

At present, there are many calculation methods for predicting drug combinations. Pal and Berlow (PAL and BERLOW, 122011) proposed a model utilizing the set theory and the information about tumor drug sensitivities and kinase inhibition profiles to predict the drug combinations. The analysis results of their model help identify drug combinations that inhibit a minimal set of kinases to decrease negative side effects. Huang et al. (Huang et al., 2014) developed a network-based sorting algorithm called DrugComboRanker which used patient genomic information and protein interaction data to construct disease-signaling networks and model a drug functional network based on drug genomic profiles. Their model final predicts potential drug combinations would be the drug-drug pairs with high overlap in the disease network, affecting multiple key signaling modules. Based on the hypothesis that drug-drug pairs with more similar chemical structures, target proteins, adverse drug reactions, and therapeutic purposes have a higher probability of effective drug combinations, Cheng and Zhao (Cheng and Zhao, 2014) applied machine learning techniques to predict drug combinations. Therefore, they calculated a lot of drug-drug similarities, such as phenotypic similarity, therapeutic similarity, chemical structure similarity, and genomic similarity. Finally, they tested the model on antipsychotic drug-drug interactions. Chen *et al.* (Chen et al., 2016) developed a model named Network-based Laplacian regularized Least Square Synergistic drug combination prediction (NLLSS) to predict potential synergistic drug combinations. They also carried out biological experiments, finally, 7 of the 13 antifungal synergistic drug combinations predicted by the NLLSS were verified. Ligeti et al. proposed a predictive model by applying a propagation algorithm on a protein-protein association network (Ligeti et al., 2013 - 2013), but the construction of the network is hampered by the incompleteness of the biological data.

However, the existing prediction methods use data from two-drug combinations, and many drug combinations are more than two drugs. Generally speaking, for the predicting efficacious drug combination of the two drugs, we can construct a graph model to score the candidate drug-drug pairs. Unfortunately, simple graphs can only model pair-wise relationships between vertexes, which prevents us from capture high-order relations. Constructing the data into a graph will lose a lot of useful information, which is not conducive to the final prediction results. In the graph theory, hypergraphs are generalizations of simple graphs where the edges are called hyperedges and the hyperedges are any subset of vertexes. Briefly, a hyperedge can have more than two vertices, that is, in this paper, it can be said that a drug combination can have more than two drugs. Therefore, it is significant to model data into hypergraphs for the utilization of data information.

In this paper, we constructed the drug combination data as a hypergraph and predicted the efficacious drug combination using the random walk with restart algorithm on the hypergraph. As a result, HRWR achieved reliable predictions with cross-validation and it obtains higher AUROC than other methods. Case studies of breast cancer, colorectal cancer, and lung cancer demonstrated the effectiveness and potential value of HRWR in identifying novel drug combinations.

## Materials and methods

### 2.1 Data collection and pre-processing

The Drug Combination Database (DCDB) was the first specialized database to collect and organize drug combination information, which includes 1363 drug combinations, consisting of 1735 drugs in all. Besides, DCDB collected 237 non-efficacious drug combinations, which may provide valuable data support for finding efficacious drug combinations. In this paper, we set efficacious drug-drug interactions as positive samples and set non-efficacious drug-drug interactions as negative samples. Drug combinations corresponding to lung cancer, breast cancer, and colon cancer in DCDB are the data set examined in this paper. In detail, the data in the database is selected according to the following criteria: ICD10 is an internationally unified disease classification method developed by WHO (Quan et al., 2005), each drug combination of the DCDB database had at least one ICD10 code. If the ICD10 Code of a drug combination is C18, i.e., Malignant neoplasm of colon, we record this data as lung cancer data. Similarly, the drug combinations which the ICD10 Code is C50 and C34, which are recorded as breast cancer and lung cancer data, respectively. Also, we did not use those combinations whose effect type were “Need further study” in the DCDB, because the experimental results of these drug combinations were still unclear.

Some drug combinations have two different types, namely efficacious and non-efficacious. In this case, we consider this combination of drugs to be efficacious. The specific reasons are as follows: when the dosage of a drug in the drug combination is too large, the drug combination will be judged to be non-efficacious because of the increase of drug side effects and toxicity. Therefore, if two drugs can be efficaciously combined once, then we can find a reasonable dose ratio to make the combination efficacious, which can be used in clinical practice.

The basic attributes of the drug combination network are shown in **Table 1**. Here, the hyperedge represented efficacious drug combination, the vertex represented the drug involved in the efficacious drug combinations. In particular, 2-edge represented an efficacious drug combination of two drugs. The non-efficacious drug combinations are the total non-efficacious drug combinations associated with the drugs involved in the efficacious drug combinations. For example, the symbol *x(y)* represents a total of *x* non-efficacious drug combinations, where *y* drug combinations in *x* drug combinations are combinations of two drugs.

**Table. 1.**
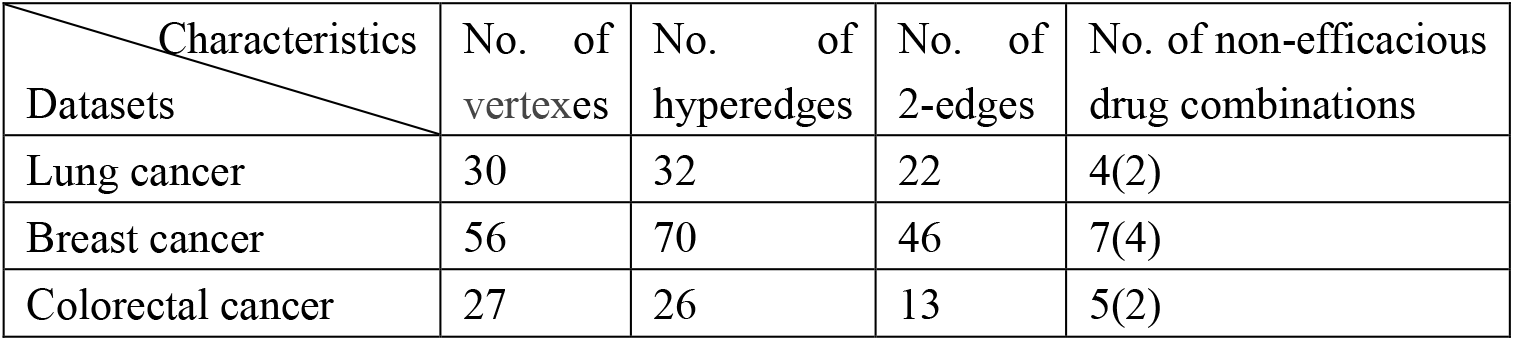
Global characteristic of the drug combination network.

### 2.2 Notations and definitions

By modeling the drug combination data as a graph, we capture the relationship between drugs where we set drugs as vertices and the combination relationships as edges.

Let *HG(V,E)* be a hypergraph with the vertex set *V* and the hyperedges set *E*. A hyperedge *E* is a subset of *V* where U_e∈E_ *e* = *V*. The hypergraph is said to be connected when there is a path between each pair of vertices. A path is a sequence of vertices over hyperedges {*v*_1_, *e*_1_, *v*_2_, *e*_2_,…,*e*_*k*-1_,*v*_*k*_} where {*v*_*i*_, *v*_*i*+1_} ⊆ *e_i_*. A hyperedge *e* is said to be incident with *v* when *v ∈ e.* A hypergraph has an incidence matrix *H* ∈ ℝ^|*V*|×|*E*|^ as follows:

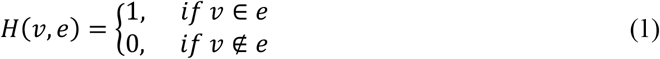

The vertex and hyperedge degree are defined as follows:

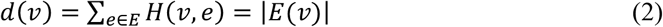

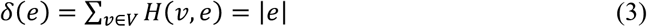

*D_e_* ∈ ℝ^|*E*|×|*E*|^ and *D_v_* ∈ ℝ^|*V*|×|*V*|^ are the diagonal matrices representing the degrees of hyperedges and vertices, respectively.

### 2.3 Random walk on hypergraph

Random walk on the simple graph has been studied extensively in many important tasks, such as ranking (Bubnicki and Orski, 2005; Minkov and Cohen, 2007), similarity and recommendation (Tayebi et al., 2011; Li et al., 2012), or link prediction (Backstrom and Leskovec, 2011). The random walk on the hypergraph is a generalization of it on the simple graph. Therefore, the random walk on a hypergraph is a transition process of the hypergraph as follows: the current vertex selects a hyperedge associated with the current vertex, and if the current vertex has more than one hyperedge associated with it, the selection is based on the hyperedge weight, i.e., the greater the weight, the greater the probability of selection. Then the current hyperedge randomly chooses a neighbor vertex within the current hyperedge as the next vertex of the transition process. Briefly, each step of the transition process is from one vertex to another. Repeat the step until the convergence condition is reached.

Consequently, we also regard the vertex set {*v*_1_, *v*_2_,…,*v_n_*} as a set of states {*s*_1_,*s*_2_,…,*s_n_*} in a finite Markov chain ℳ. The transition probability of ℳ is a conditional probability defined as *P(u,v) = Prob(s_t+1_ = v|s_t_ = u)* which means that the ℳ will be at *v* at time *t* + 1 given that it was at *u* at time *t*. What is more, for any *u* of *V* we have *∑_v∈V_P(u,v)* = 1.

Define transition probability *P* ∈ ℝ^|V|×|V|^ as follows:

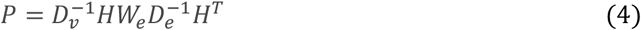

Define *r_t_* ∈ ℝ^|V|×1^ as a vector in which the *i*-th element represents the probability of discovering the random walk at vertex *i* at step *t*, so the probability *r_t+1_* can be calculated iteratively by:

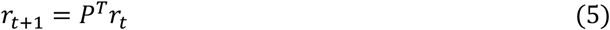

For the random walk with restart algorithm (Tong et al., 2006 - 2006), there is an additional restart item compared to the above algorithm. The probability *r_t+1_* can be calculated iteratively by the following expression:

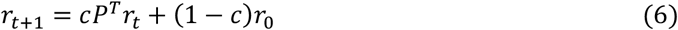

Define initial probability *r_0_* ∈ ℝ^|V|×1^ as a vector in which the *i*-th element is equal to one, while other elements are zeros. And *1 - c* is the restart probability (0 ≤ c ≤ 1).

The random walk will be stable when the *L1* norm between *r_t+1_* and *r_t_* is less than a cutoff. Here we took the cutoff as 10^-6^, and this criterion is often used in some researches (Köhler et al., 2008; Chen et al., 2010; Li and Patra, 2010; Chen et al., 2012). When the is stable, stable probability between vertex *i* and vertex *j* is defined as the *j*-th element of *r_t_* corresponding to the starting vertex is *v_i_*. **Fig. 1** is an example of the random walk on a hypergraph.

**Fig 1.**
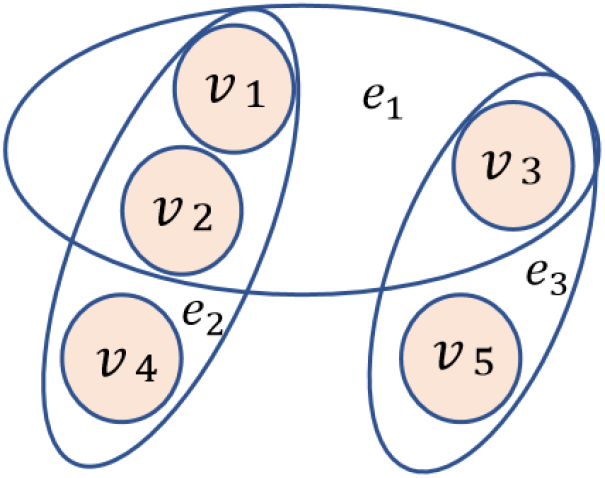
An example of the random walk on a hypergraph. First, Suppose the initial vertex is v_1_, it has *1/2* probability to choose *e_1_* and *1/2* probability to choose *e_2_.* Then, if *e_2_* is selected, *v_1_* has *1/3* probability to choose *v*_1_, *v_2_,* or *v_4_* as the next vertex of the transition process. Next, repeat the process of “select hyperedge first, then select vertex”. Finally, when the number of random walks is enough (t tends to infinity), we can get the “selected” probability of each vertex, that is, the steady-state probability.

## Results

In this section, cross-validation evaluated the predictive performance of HRWR first. Then, based on the golden standard datasets, we also conducted case studies to verify the efficiency of HRWR in discovering potential combinatorial drugs.

### 3.1 Cross-validation tests

A receiver operating characteristic (ROC) curve is a graphical plot that shows the diagnostic ability of the binary classifier system because its recognition thresholds are different. AUROC (Area Under Receiver Operating Characteristic Curve) is the area under the AUROC curve with a value between 0 and 1. AUROC can intuitively evaluate the quality of the classifier, the larger the value, the better.

According to the different types of cross-validation, the drug pairs corresponding to the elements of 1 in the adjacent matrix A are proportionally used as the training set and the test set, and the efficacious score between each pair of drugs is predicted.

In this paper, we used two different types of cross-validation for different methods and data sets to test the predictive ability of the model, we did the following two types of cross-validation:

1. Compare the value of LOOCV when the hyperedge weight is proportional to and inversely proportional to the number of vertices contained in the hyperedge. As shown in **Fig. 2**, changing the weight of hyperedge shows that the effect is better when the hyperedge weight is proportional to the number of vertices contained by hyperedge.
2. Remove a vertex-vertex pair randomly. In this case, HRWR is compared with the random walk with restart (RWRDC) (Wang and Yan, 2020), random walk, and NLLSS (Chen et al., 2016). To compare the predictive power of different models more equitably, we all use the same data set, that is, we only use the data of existing drug combinations, but not the data of drug similarity. Especially, we regarded each drug as the main drug and the adjuvant drug for the NLLSS model.

**Fig. 2.**
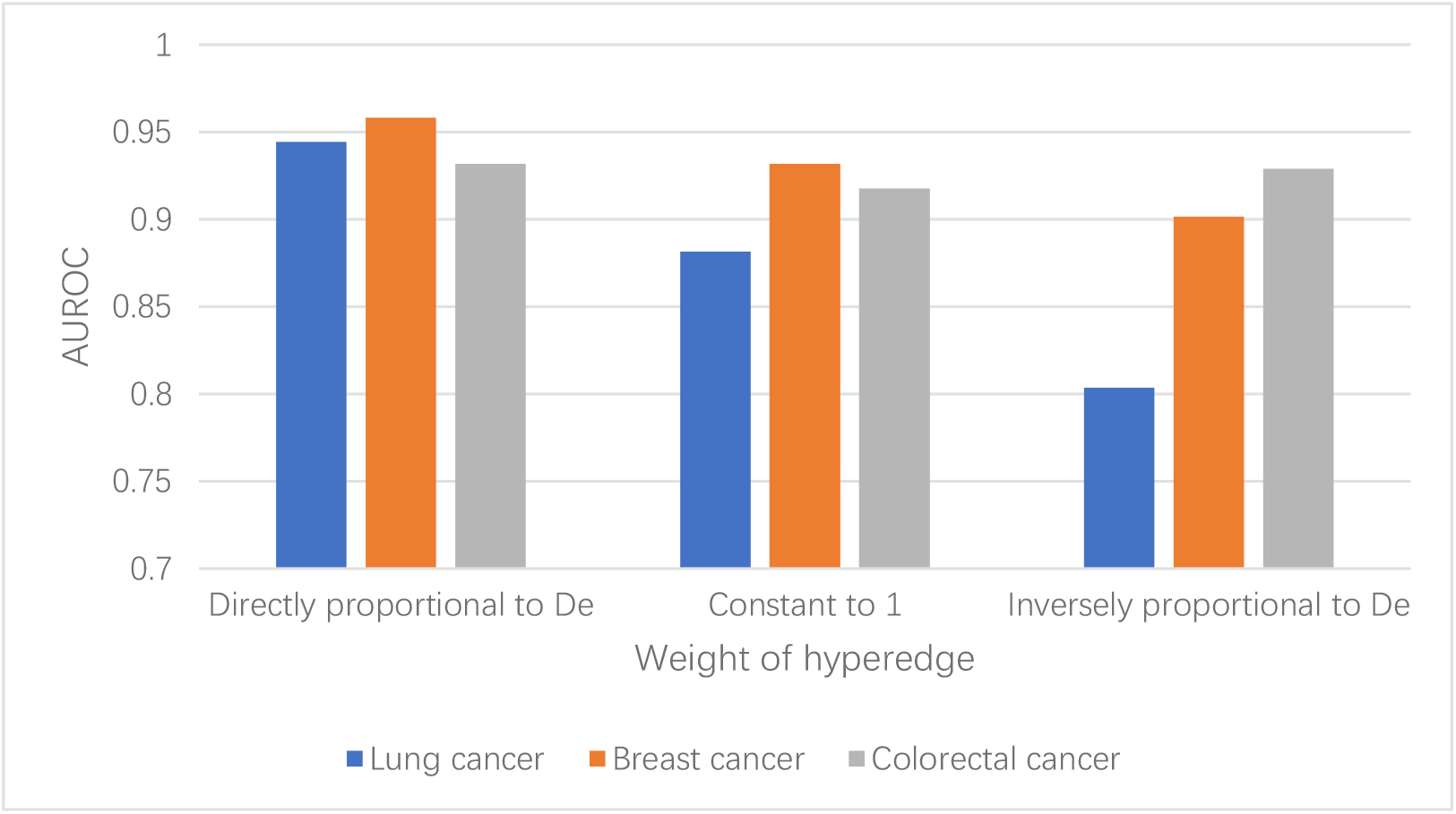
The performance of different weight of hyperedge on the LOOCV.

As shown in **Fig. 3**, in the case of LOOCV, the blue line represented the ROC curve of HRWR, the green line, dark yellow line, and purple line represented the ROC curve of the RWRDC, random walk model, and NLLSS, respectively. The graphs show that the prediction result of HRWR is better than others.

**Fig. 3.**
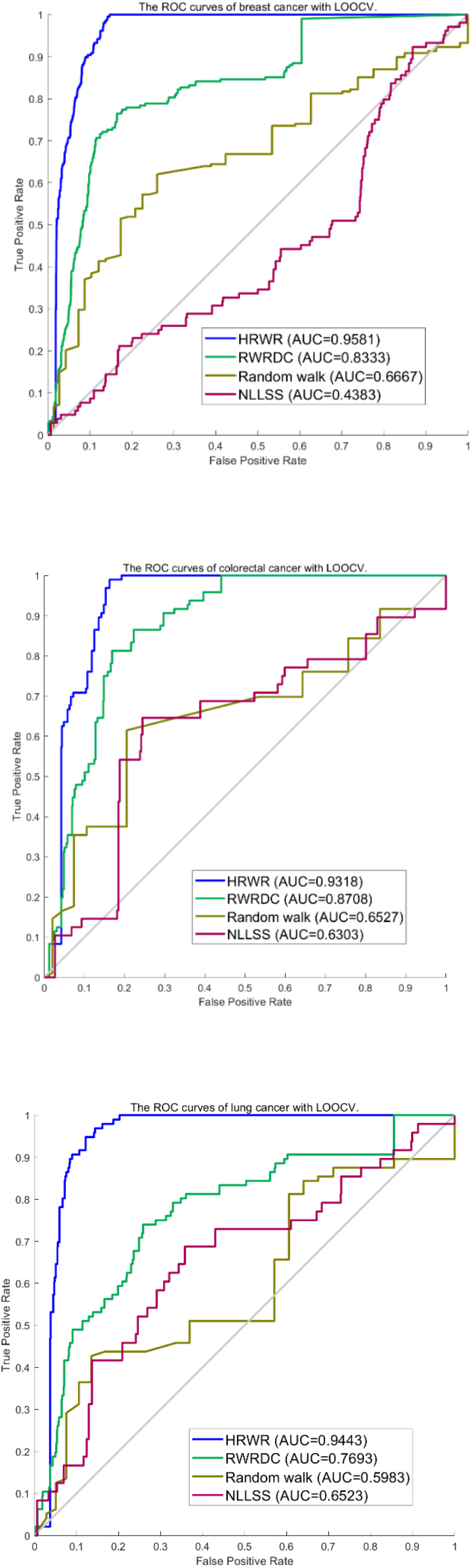
The ROC curves of breast cancer, colorectal cancer, and lung cancer with LOOCV.

### 3.2 Case studies

The case study aimed to examine the capability of HRWR in discovering novel efficacious drug combinations. For lung cancer, breast cancer, and colorectal cancer, we ranked all drug-drug pairs based on their corresponding predicted efficacious scores. Prediction results were verified based on not only the DCDB database but recently published experimental literature. **Table 1** shows that efficacious drug combinations account for about 5%-9% of the total combinations of each cancer data set, therefore, we analyzed the top 10% predictions. **Table 2-4** listed all the corresponding top 10% ranking results. The “Null” in the table indicates that documentation has not been found.

**Table. 2.**
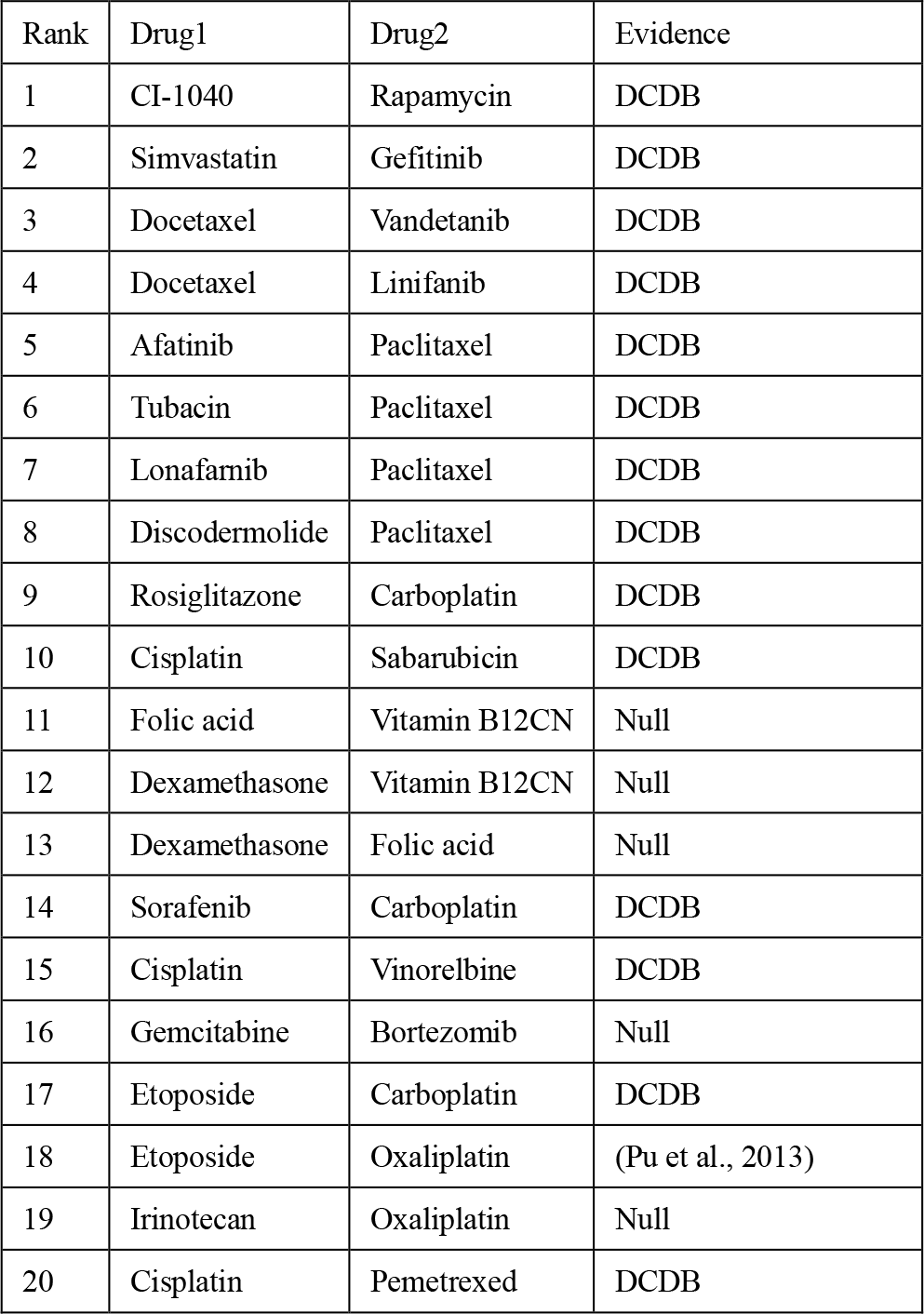
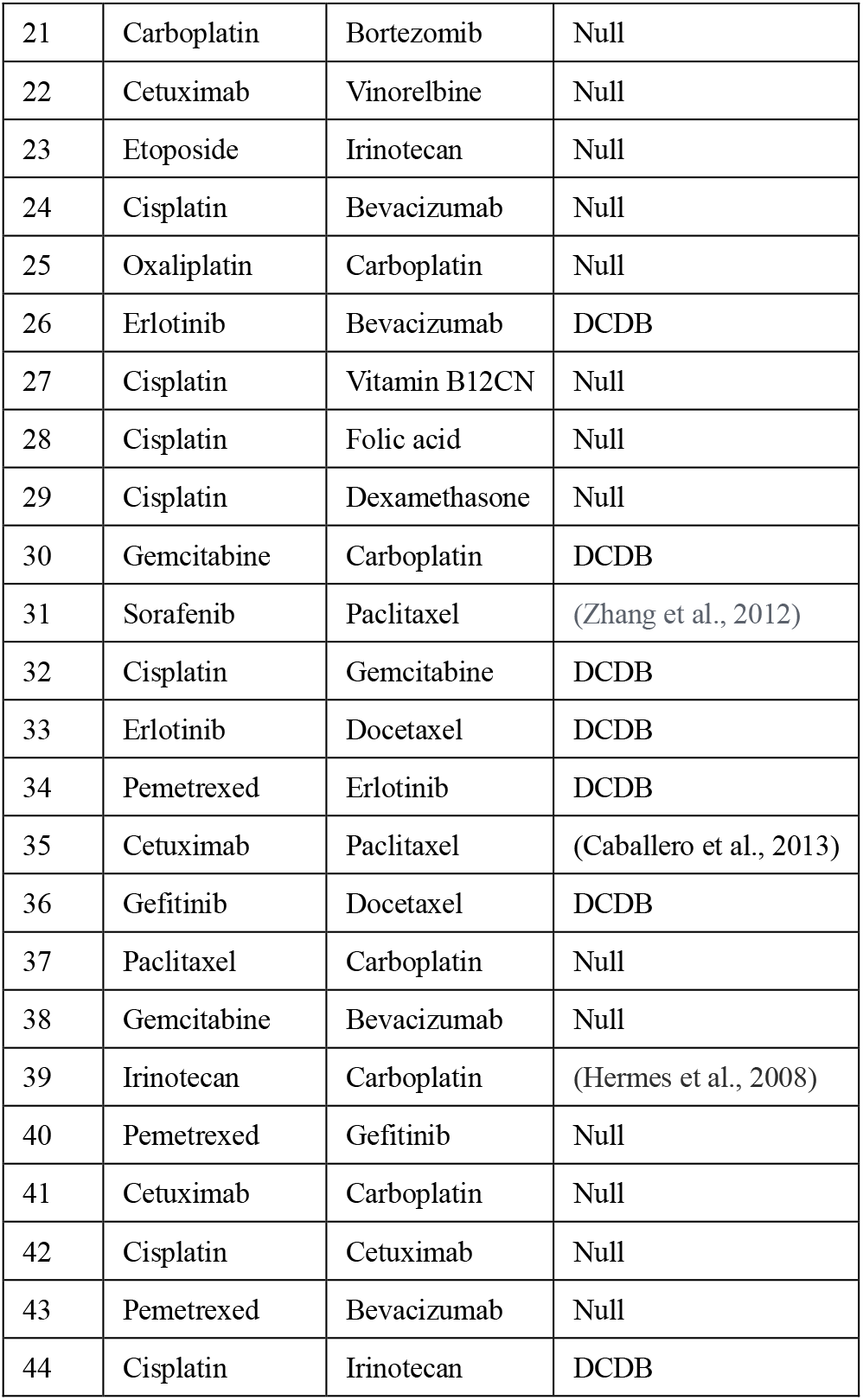
Case study of lung cancer.

Lung cancer originates in the tissues of the lungs or the cells lining the airways, the bronchi. The disease begins with mutations of normal cells that turn them into cancer cells. These cells divide and multiply at an out-of-control rate and can eventually form tumors that impede breathing and oxygenation throughout the body (Dela Cruz et al., 2011). We used HRWR to predict potential efficacious drug combinations. As a result, all potential lung cancer related efficacious drug combinations that ranked top 10 had been validated by databases and recent experimental literature. When we took the drug-drug pairs of top 10% the highest predicted efficacious score as the predicted result, all the known efficacious drug combinations contained in the data set would be successfully predicted. What is more, Etoposide-Oxaliplatin, Sorafenib-Paclitaxel, Cetuximab-Paclitaxel, and Irinotecan-Carboplatin were predicted to be 18^th^, 31^th^, 35^th^, and 39^th^ by HRWR respectively. These efficacious drug combinations are not recorded in the database of DCDB, but we have verified the effectiveness of drug combinations through the latest published literature.

Breast cancer is the most frequently diagnosed cancer and the leading cause of cancer-related mortality in females worldwide (Zheng et al., 2014), which comprises 22% of all cancers in women (Donahue and Genetos, 2013; Karagoz et al., 2015). 70 groups of efficacious drug combinations have been discovered by biological experiments, HRWR can also predict more potential efficacious drug combinations. Consequently, 100% of drug-drug pairs that ranked top 10 in the data set had been validated, respectively. When we took the drug-drug pairs of top 10% the highest predicted efficacious score as the predicted result, all the known efficacious drug combinations contained in the data set would be successfully predicted. Besides, 22 groups of efficacious drug combinations have been validated by the latest clinical trial literature (see **Table 3** for details). Although some predictions have not been validated in databases and literature, there is still a strong possibility that predicted efficacious drug combinations will be available. For example, Irinotecan-Oxaliplatin was predicted to be 19^th^ by HRWR and there was no direct evidence that it was an efficacious drug combination. But (Lombard et al., 1991) showed that the regimen of irinotecan plus oxaliplatin has higher efficacy and less myelosuppression as compared to etoposide plus oxaliplatin in the treatment of extensive-stage SCLC. Although there is a higher incidence of diarrhea, it can be tolerant.

**Table. 3.**
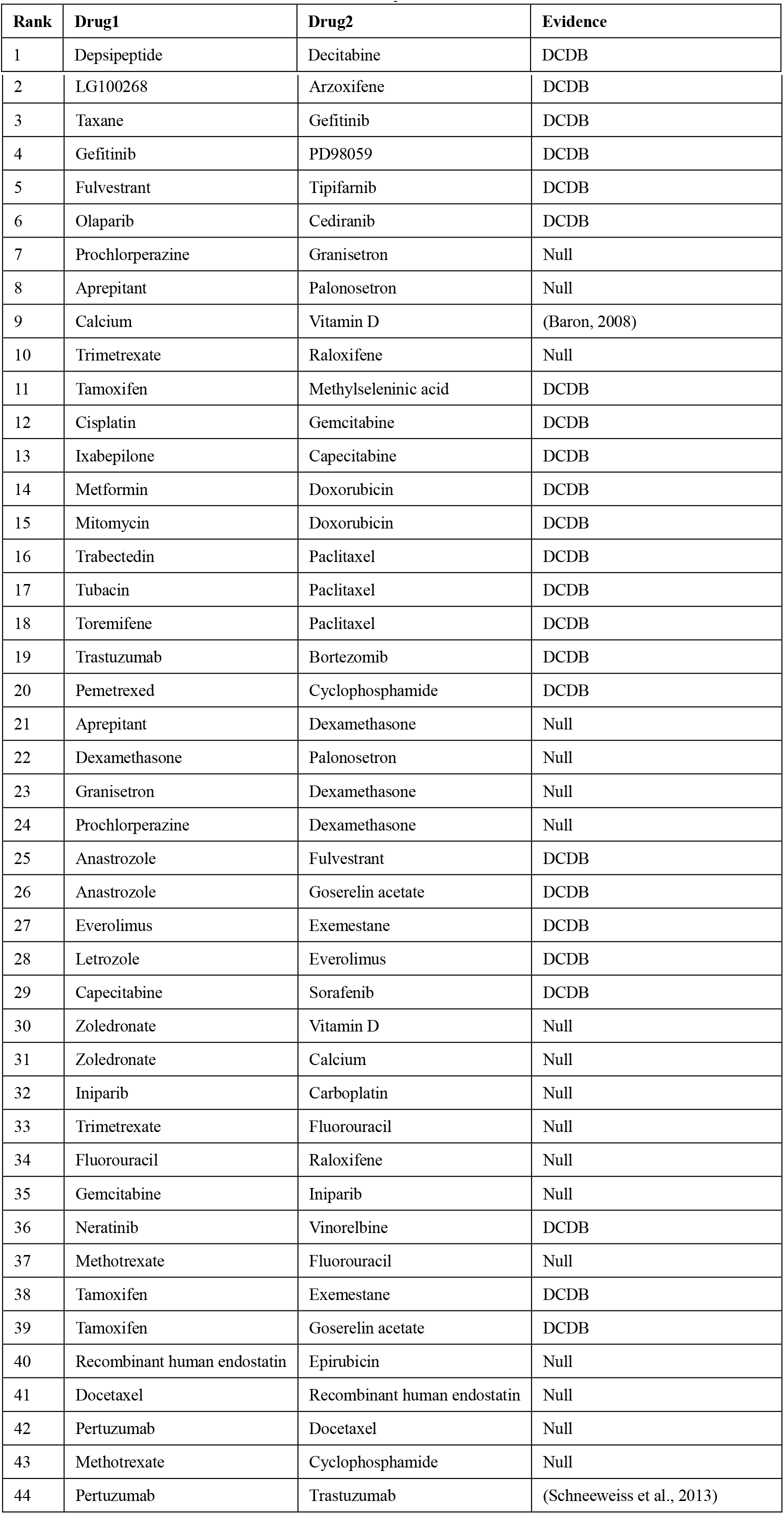
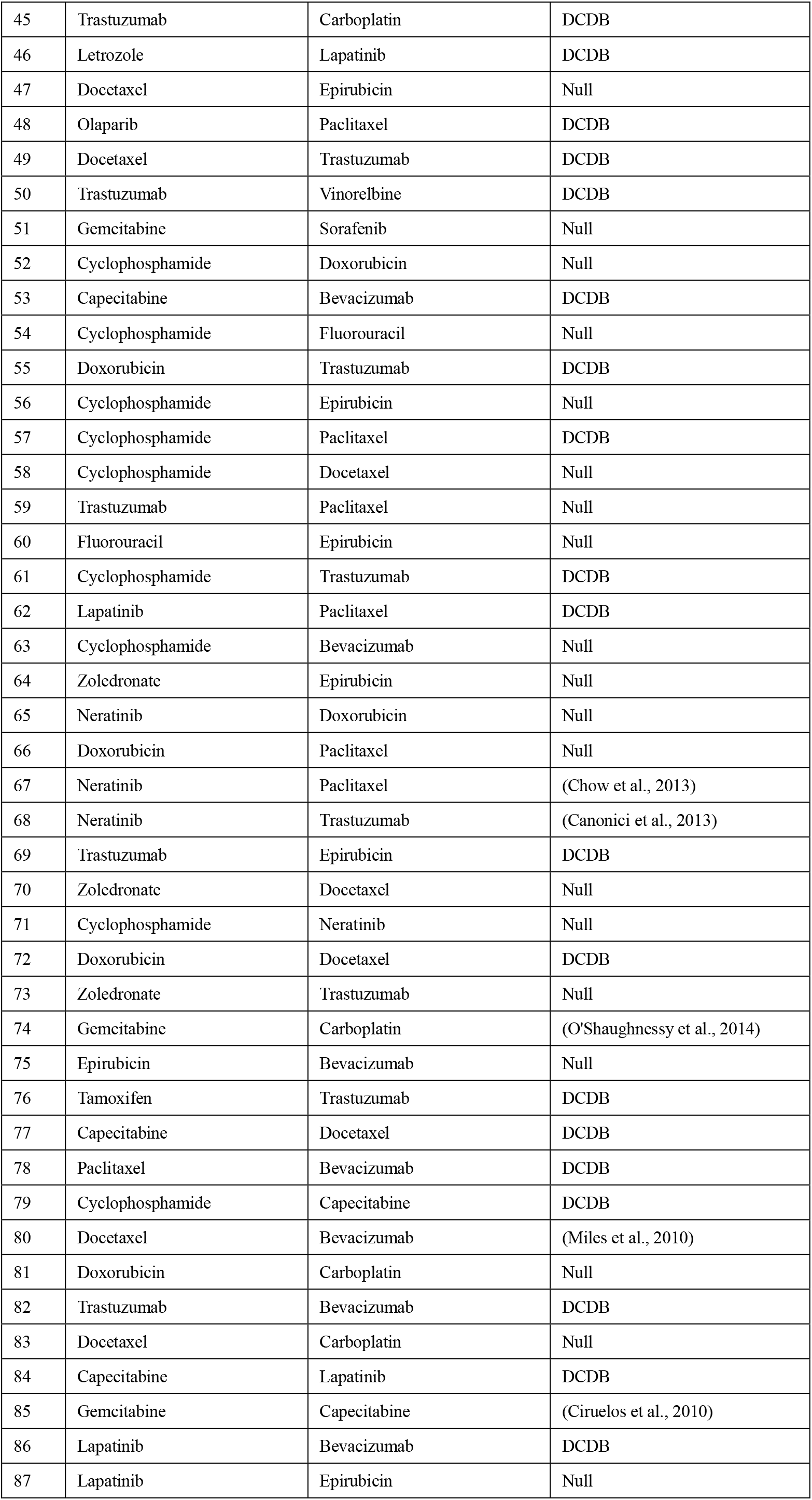
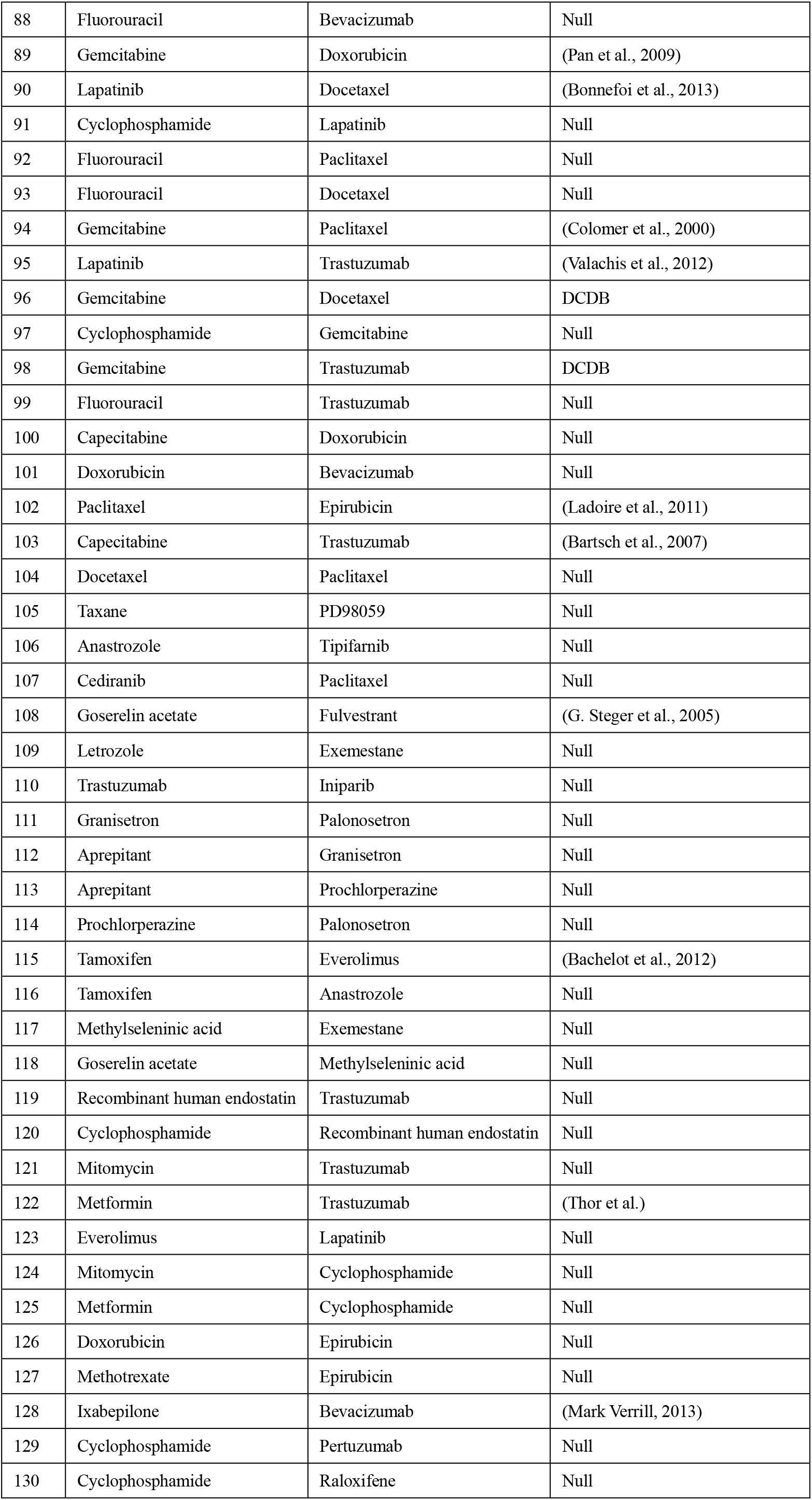
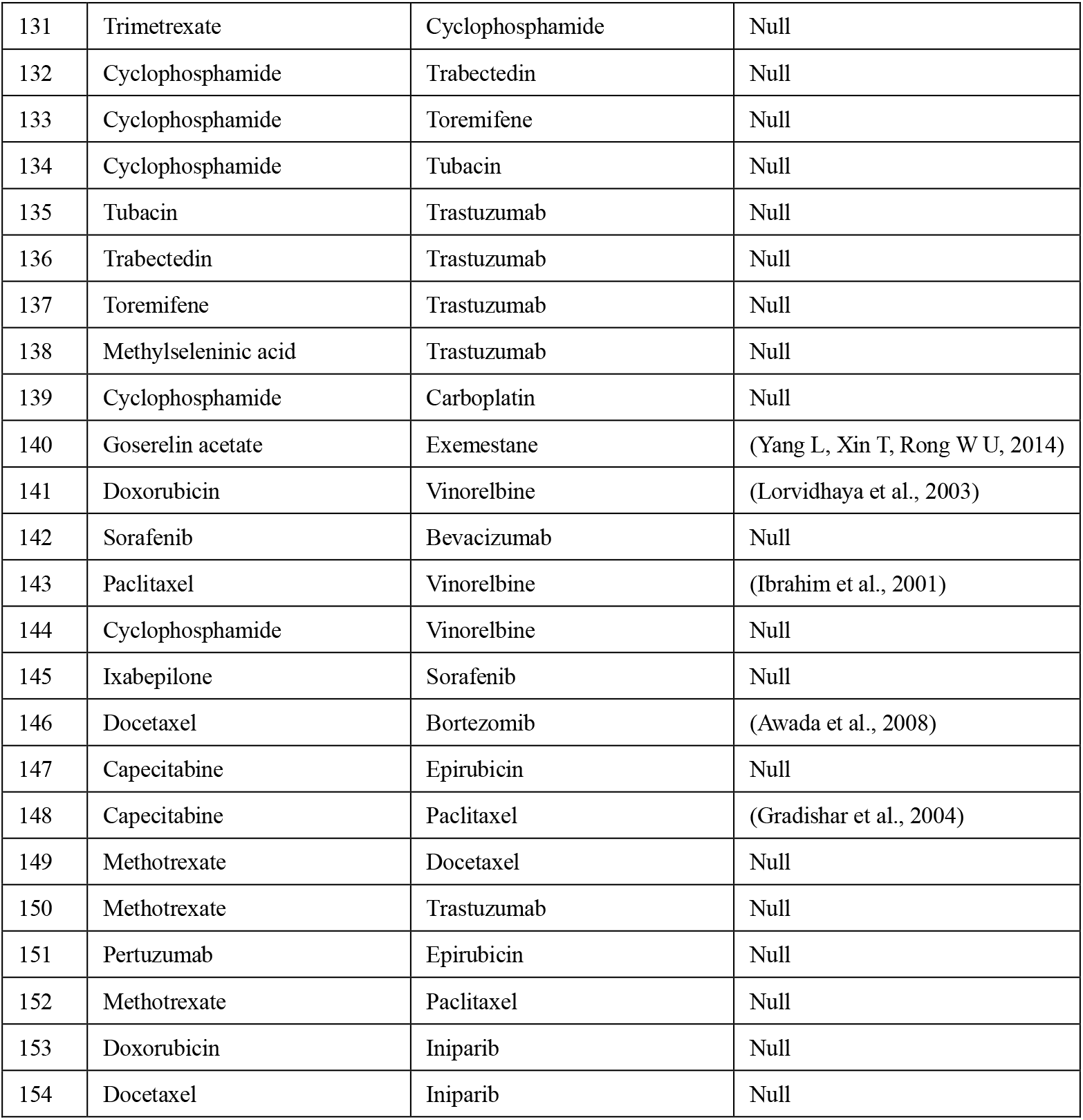
Case study of breast cancer.

Colon cancer is one of the most common malignant tumors in the world (Xue et al., 2015), killing almost seven hundred thousand people every year (Gu et al., 2017), even the disease-specific mortality rate is close to 33% in the developed countries (Han et al., 2015). From **Table 4**, we could know that all efficacious drug combinations in the data set had been predicted by HRWR. Besides, the combinations of Fluorouracil and Bevacizumab, Cetuximab and Capecitabine, Cetuximab and Bevacizumab were predicted to be 29^th^, 34^th^, and 35^th^ by HRWR respectively. These efficacious drug combinations are not recorded in the database of DCDB, but we have verified the effectiveness of drug combinations through the latest published literature. These results showed that HRWR could predict more potential efficacious drug combinations.

**Table. 4.** Case study of colorectal cancer.

| Rank | Drug1 | Drug2 | Evidence |
| --- | --- | --- | --- |
| 1 | Calcium carbonate | Cholecalciferol | DCDB |
| 2 | Temozolomide | Lomeguatrib | DCDB |
| 3 | Propranolol | Etodolac | DCDB |
| 4 | Trifluridine | Docetaxel | DCDB |
| 5 | Irinotecan | Gefitinib | DCDB |
| 6 | Fluorouracil | RPR-115135 | DCDB |
| 7 | Fluorouracil | 2'-Deoxyinosine | DCDB |
| 8 | Dasatinib | Oxaliplatin | DCDB |
| 9 | 5-2,4-dihydroxypyrimidine | Tegafur | Null |
| 10 | Potassium oxonate | Tegafur | Null |
| 11 | Potassium oxonate | 5-2,4-dihydroxypyrimidine | Null |
| 12 | Dexamethasone | Floxuridine | Null |
| 13 | Leucovorin | Fluorouracil | Null |
| 14 | Capecitabine | Oxaliplatin | DCDB |
| 15 | Leucovorin | Methotrexate | Null |
| 16 | Methotrexate | Fluorouracil | Null |
| 17 | Floxuridine | Capecitabine | Null |
| 18 | Dexamethasone | Capecitabine | Null |
| 19 | Irinotecan | Fluorouracil | Null |
| 20 | Leucovorin | Irinotecan | Null |
| 21 | Leucovorin | Panitumumab | Null |
| 22 | Floxuridine | Oxaliplatin | Null |
| 23 | Dexamethasone | Oxaliplatin | Null |
| 24 | Panitumumab | Fluorouracil | Null |
| 25 | SC144 | Fluorouracil | DCDB |
| 26 | SC144 | Oxaliplatin | DCDB |
| 27 | Panitumumab | Oxaliplatin | Null |
| 28 | Irinotecan | Bevacizumab | Null |
| 29 | Fluorouracil | Bevacizumab | (Kabbinavar et al., 2009) |
| 30 | Capecitabine | Bevacizumab | DCDB |
| 31 | Leucovorin | Oxaliplatin | Null |
| 32 | Leucovorin | Bevacizumab | Null |
| 33 | Irinotecan | Oxaliplatin | DCDB |
| 34 | Cetuximab | Capecitabine | (Sastre et al., 2012) |
| 35 | Cetuximab | Bevacizumab | (Nygren et al., 2005) |

## Discussion

This paper first models the drug combinations into hypergraphs. The hypergraph is a kind of high-order graph representation of biological data, it could overcome information-loss in normal graph methodology which could only describe pair-wise relationship structure. Furthermore, this is the first drug combination prediction model that uses information on more than two drugs. Due to the use of higher-order information, such as the combination of three drugs, the model has better predictive performance than the random walk on the simple graphs. HRWR can be predicted only by combining its information, that is, no additional information is needed. In this way, the process of sorting out all kinds of data is omitted when predicting potential drug combinations.

For the same disease, the therapeutic effect of a drug combination can be categorized not only as effective and non-efficacious, but also as better than monotherapy, active and safe, and so on. However, due to a large amount of missing data in the current database and the inconsistent recording in the literature, it is difficult to determine the degree of efficacy. There is no consistent way, i.e., no quantitative criteria. However, the HRWR in this paper can give validity scores for different drug pairs, and although it is not possible to verify the accuracy of such scoring, the scoring is reliable based on the only documented evidence, which is a breakthrough in drug combination modeling, and other researchers can continue to study the magnitude of effectiveness in predicting drug combinations. The results of HRWR demonstrates the power of harnessing network-based drug-drug interactions and provide additional filtering rules for drug combination studies.

Although the model performs well on the predicted results, there are still some aspects that can be improved. At present, there are few computational models for predicting multiple (i.e., more than two) efficacious drug combinations. Although the method proposed in this paper utilizes information on multiple drug combinations, it cannot predict whether a combination of more than two drugs is efficacious. Thus, predicting efficacious multiple drug combinations could be future research work. Moreover, HRWR only uses the information on the drug combination itself and does not introduce more interactive information, such as the drug-target interactions, drug-pathway interactions, disease-gene associations, and side-effects of drug combinations. Besides, the choice of restart probability has not been combined with the biological background of the drug combination, and it would make sense to give biological significance to the probability of restart in the future. What is more, HRWR could not predict interactions for novel drugs for which no interaction information is currently available.

## Declarations

## Ethics approval and consent to participate

Not applicable.

## Consent to publish

Not applicable.

## Availability of data and materials

Publicly available datasets were analyzed in this study. This data can be found here: http://www.cls.ziu.edu.cn/dcdb/. (Temporary DCDB database address: http://public.synergylab.cn/dcdb/.)

The code and dataset of HRWR are freely available at https://github.com/wangqi27/HRWR.

## Authors’ Contributions

QW conceived the project, developed the prediction method, designed the experiments, implemented the experiments, analyzed the result, and wrote the paper. GY conceived the project, analyzed the result, and revised the paper.

All authors have read and approved the manuscript.

## Acknowledgements

This work was supported by the National Natural Science Foundation of China and the National Key Research & Development Program of China. We thank Dr. Ming Chen *et al.* for making the drug combination data complete, published in ref. (Liu et al., 2014), freely available via the DCDB Database.

## Authors’ Information

### Affiliations

Academy of Mathematics and Systems Science, Chinese Academy of Sciences, Beijing, 100190, China.

Qi Wang, Guiying Yan

University of Chinese Academy of Sciences, Beijing, 100190, China.

Qi Wang, Guiying Yan

### Corresponding author

Correspondence to Guiying Yan.

